# Sea squirt-inspired bio-derived tissue sealants

**DOI:** 10.1101/2023.10.02.560578

**Authors:** Aishwarya V. Menon, Jessica E. Torres, Abigail D. Cox, Marije Risselada, Gudrun Schmidt, Jonathan J. Wilker, Julie C. Liu

**Affiliations:** Davidson School of Chemical Engineering, Purdue University, West Lafayette, IN 47907, USA; Department of Chemistry, Purdue University, West Lafayette, IN 47907, USA; Department of Comparative Pathobiology, Purdue University, West Lafayette, IN 47907, USA; Department of Veterinary Clinical Sciences, Purdue University, West Lafayette, IN 47907, USA; School of Materials Engineering, Purdue University, West Lafayette, IN 47907, USA; Weldon School of Biomedical Engineering, Purdue University, West Lafayette, IN 47907, USA

**Keywords:** biomedical glue, burst pressure, gallol-based glue, nature-inspired glue, tunicate- mimetic glue

## Abstract

Sea squirts’ or tunicates’ bodies are composed of cellulose nanofibers and gallol- functionalized proteins. These sea creatures are known to heal their injuries under seawater by forming crosslinks between gallols and functional groups from other proteins in their bodies. Inspired by their wound healing mechanism, herein, we have developed a tissue sealant using zein (a plant-based protein) and tannic acid (gallol-containing polyphenol). Except for fibrin-based sealants, most commercial surgical adhesives, and sealants available today are derived from petroleum products that compromise their biodegradability. They often have complicated and multi-step synthesis processes that ultimately affect their affordability. To overcome this challenge, we ensured that these sea squirt-inspired tissue sealants are bio-based, easily synthesized, and low-cost. The sealants were studied on their own and with a food-grade enzyme transglutaminase. The adhesion performances of the sealants were found to be higher than physiological pressures in seven out of nine different tissue substrates studied here. Their performance was also better than or on par with the FDA-approved fibrin sealant Tisseel. *Ex vivo* models demonstrate instant sealing of leaking wounds in less than a minute. The sealants were not only cytocompatible but also showed complete wound healing on par with sutures and Tisseel when applied *in vivo* on skin incisions in rats. Overall, these sea squirt-inspired bio-based sealants show great potential to replace currently available wound closure methods.

## Introduction

Achieving complete wound closure is a big challenge in most surgeries around the world. The drawbacks of traditional mechanical wound closure methods (e.g., sutures and staples) have led to the emergence of non-invasive wound closure methods such as surgical tissue adhesives and sealants.[1–4] One of the first adhesives to be commercialized for surgical applications were cyanoacrylates (e.g., butyl and octyl cyanoacrylate) and fibrin-based sealants. Cyanoacrylates have been in surgical use for over 50 years, whereas fibrin sealants have been the most widely used surgical adhesive in the U.S. since 1998 after their first approval by the FDA.[5] Cyanoacrylate adhesives (e.g., Trufill, Omnex) exhibit fast gelation times and instant tissue-sealing properties. However, cyanoacrylates are stiffer than soft tissues, and they break down within the body to toxic byproducts including formaldehyde. As a result, they are used strictly for topical adhesion applications only.[6–8] Fibrin sealants (e.g., Tisseel, Vistaseal) are two component sealants consisting of fibrinogen and thrombin. When applied to a site, the components react together to mimic a blood clot and seal the wounds. Though these turned out to be successful initially, they exhibit poor adhesion strengths and pose the risk of thrombosis, embolism, and transmission of serological diseases due to their source being pooled donor blood.[9]

These limitations of the earliest commercially available tissue sealant have led to the emergence of newer sealants based on other types of wet-adhesion chemistries such as albumin and glutaraldehyde (e.g., BioGlue, PreveLeak), *N*-hydroxysuccinimide (NHS) reactive esters (e.g., Progel, Coseal), and polyurethanes (e.g. TissuGlu).[2,4] Although these new sealants have shown immense promise, there is still a need for better alternatives not only due to toxicity and poor adhesion concerns but also because most of them are of petroleum origin which causes biodegradability issues. Moreover, they also often involve tedious synthesis routes and can be expensive and unprofitable.

Bioinspired materials have gained a lot of traction in recent years. In this regard, adhesives inspired by sea creatures such as mussels and barnacles are being increasingly researched.[10–16] Tunicate or sea squirt is one such marine organism whose wound-healing mechanism underwater has inspired the development of wet adhesives.[17–19] The adhesion mechanism of mussels is based on the catechol-containing amino acid 3,4-dihydroxyphenylalanine (DOPA) whereas the wound healing mechanism in sea squirts is based on the gallol-containing amino acid called 3,4,5-dihydroxyphenylalanine (TOPA).[20] Sea squirt bodies are composed of cellulose nanofiber and gallol-functionalized peptides. They heal injuries under seawater using gallol chemistry to bond to functional groups in proteins (figure 1a). Additionally, gallols in TOPA are also known to form coordination complexes with metal ions, that again act as an adhesive for healing injuries in sea squirts.[18]

**Figure 1:**
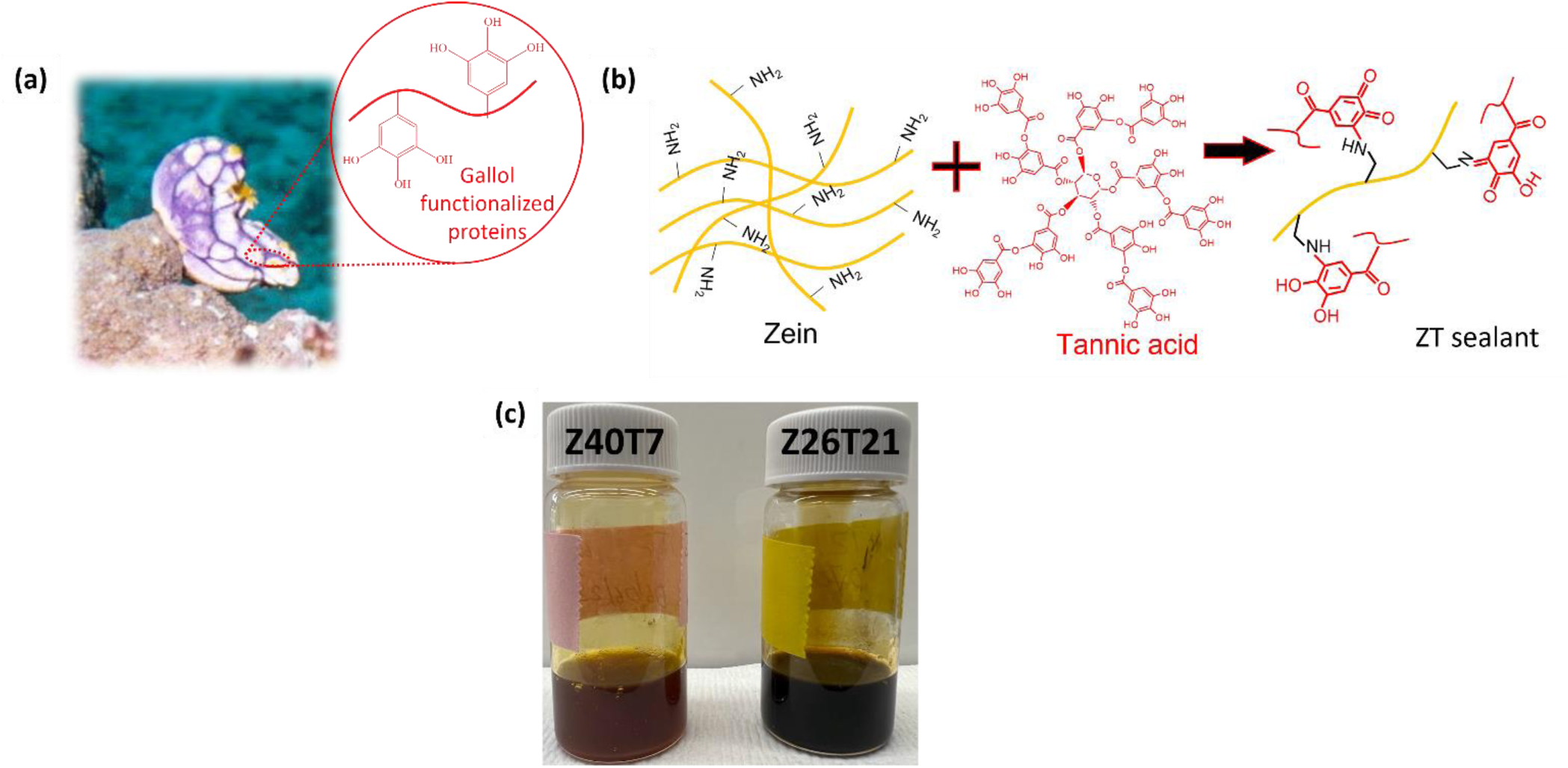
**(a**) In the ocean, sea squirts use gallol chemistry to crosslink to proteins and metal ions for wound healing. **(b)** The crosslinking mechanism of sea squirt-inspired ZT sealants. **(c)** Optimized ZT sealant formulations are darker in color with increasing TA concentration. Z40T7 is lighter in color compared to Z26T21, which has a higher TA concentration.

We recently developed adhesives from zein corn protein and gallol-containing tannic acid (TA). These glues were as strong as commercial cyanoacrylate (e.g., Super Glue) when measured under similar conditions on aluminum substrates.[21–23] When cured in wet conditions (e.g., seawater, saline, tap water, deionized water) the bond strength was found to increase over time.[24] These promising results encouraged us to study the potential use of these bio-based sea squirt-inspired adhesives as surgical tissue sealants. A commercial food-grade enzyme transglutaminase (TG) was also used in combination with these zein and tannic acid-based sealants (ZT sealant) to study potential improvements in burst pressure from additional crosslinking with tissues. ZT sealants were tested on nine different tissue substrates (or organs), and they could withstand pressures higher than normal physiological pressures experienced in seven out of nine of these organs. The sealant performance was better than or on par with the performance of the commercial gold standard, Tisseel. We believe that these ZT tissue sealants are among the cheapest, most sustainable, and most scalable tissue sealants reported in the literature.

## Materials and Methods

### Materials

Zein (Molecular weight:19-22kDa) and tannic acid were purchased from Sigma-Aldrich. Ethanol (190 proof) was purchased from Decon Laboratories. RM Transglutaminase (Moo Gloo™) was purchased from modernistpantry.com. All other reagents used were purchased from Fischer Scientific unless stated otherwise. All the porcine tissues and organs used in this study were either procured locally or from Animal Technologies, Inc. The FDA-approved surgical sealant, Tisseel, which was used for benchmarking, was donated by Baxter.

### Synthesis of Sea Squirt-Inspired Tissue Sealants

To determine the formulations with the highest adhesive strengths, the total concentration of zein was kept constant at 40 wt%, 30 wt%, or 26 wt%. Tannic acid concentrations were varied from 1 wt% to 28 wt%. A 3:1 ethanol-to-water ratio was used as the solvent to dissolve zein and tannic acid. To prepare the sealants, the components were combined manually using a wooden spatula. The pH of the prepared sealant was adjusted to 7.0 using a 10 M NaOH solution. The sealant was then incubated at 37 °C for 24 h to remove any bubbles before testing. Initially, the sealant was amber-colored, but the color intensified to dark brown after 24 h depending on the amount of tannic acid added.

### Burst Pressure Test

The burst pressure of the sealants was tested on porcine tissue substrates using an in-house developed burst pressure test setup based on a modified version of ASTM standard F2392 – 04.[25] The tissue substrates were cut into dimensions of approximately 2.5 cm * 2.5 cm. A 3 mm diameter puncture was made at the center of the tissue substrates using a biopsy punch. The sealant was applied to the puncture and allowed to cure before being loaded onto the test fixture. A syringe pump was used to pump test fluid into the test fixture at a rate of 5 mL/min. A pressure transducer (PX409-100GUSBH, OMEGA Engineering) connected to a computer was used to record the burst pressures.

Initial optimization studies to determine the best sealant formulations were performed on porcine skin substrates using 20 µL of the sealant. When the effect of TG was studied, 20 µL of 300 mg/mL TG was applied to the tissues followed by 20 µL of the sealant. The burst pressure increased as the sealant volume increased. Hence, after identifying the best formulations, 40 µL of the sealant and/or 40 µL of TG were applied to study the burst pressure strengths on all tissue substrates. The sealants were cured for 30 min or 2 h in a humidified incubator at 37 °C. For the 30 min cure time, the samples were cured in the humidified incubator for 20 min and then allowed to equilibrate at room temperature for 10 min. For the 2 h cure time, the samples were cured in a humidified incubator for 1.5 h and then allowed to equilibrate at room temperature for 30 min. A 2 h cure time was used during initial optimization studies. The test fluid used was 1x phosphate-buffered saline (PBS) for all tissues except for the stomach and lungs in which case simulated gastric fluid and air were used respectively. Gastric fluid was prepared by mixing 1% 12M HCl solution with 0.5 g pepsin to obtain a solution with a final pH of 2.5. All the experiments were performed with at least five replicates.

### Cytocompatibility studies

NIH/3T3 fibroblasts were cultured at 37 °C and 5% CO2 in low-glucose Dulbecco’s modified Eagle’s medium (DMEM) supplemented with 10% fetal bovine serum (Lonza, Basel, Switzerland) and 1% penicillin/streptomycin antibiotic. Cells were subcultured at 60-80% confluency. The cytocompatibility of the zein-based sealants was studied by culturing the cells in sealant extracts in accordance with the ISO 10993-5 standard.[26] Briefly, 200 µl of sealants were cured in cylindrical molds in a humid environment for 72 h at 37 °C to obtain a 4.5 mm diameter and 1 mm thick cylindrical sample. The samples were then immersed in 5 mL of DMEM at 37 °C for 24 h to obtain the leachate. The leachate was studied at three different dilutions of 1x, 10x and 100x. Leachate was sterile-filtered using a 0.2-μm filter. Cells were seeded into wells of a 24-well tissue culture treated plate at 5,000 cells/cm^2^ and incubated overnight. After cells were attached, the media was replaced with 500 μL of leachate or 500 μL of fresh media for positive control wells. Cells were then incubated for 24 h.

A PrestoBlue assay (Thermo Fisher) was used to assess metabolic activity. Leachate was removed, and wells were rinsed 3 times with 200 μL of PBS+ (PBS supplemented with 0.01 wt% calcium chloride and 0.01 wt% magnesium chloride) to prevent detachment. Negative control wells were incubated with 70% ethanol for 5 min at ambient temperature. An aliquot consisting of 500 μL of fresh media and 50 μL of 10x PrestoBlue was then added to each well, and the plates were incubated for 4 h at 37°C. After incubation, aliquots were removed and placed into 96-well plates. The fluorescence of the reagent was measured using excitation at 555 nm and emission at 595 nm.

To measure viability, a LIVE/DEAD Viability/Cytotoxicity assay (Thermo Fisher) was used. Cells were rinsed 3 times with 200 μL of PBS+. Samples were stained with 2 μM calcein-acetoxymethyl and 1.5 μM ethidium homodimer in PBS+ for 10 min at room temperature and rinsed 3 times with 200 μL of PBS+. Fluorescent images were obtained with an excitation of 489 nm and emission of 508 nm for live cells and an excitation of 552 and emission of 578 nm for dead cells.

### Ex vivo studies

To demonstrate the efficacy of our sealant to plug leaking punctures, *ex vivo* experiments were performed using porcine stomach tissue and sausage casing. Briefly, the porcine stomach was filled with water colored with red dye. The stomach was then placed underwater, a hole was punctured using a 3 mm biopsy punch, and dyed water was allowed to leak through the puncture. The sealant was then applied to the leaking puncture underwater using a syringe. To ensure complete puncture closure, the stomach tissue was then transferred to a new container of clean water to test for any leakage from the sealed puncture.

Similarly, the sausage casing was cut into an appropriate length of 20 cm and connected to a water pump that circulated water continuously through the sausage casing at a pressure of approximately 30 mmHg. A 3 mm puncture was made using a biopsy punch. The sealant was applied to the puncture with slight pressure. The water pressure was monitored during the experiment to ensure complete puncture closure.

### *In vivo* wound healing in a rat model

The *in vivo* wound closure ability of our ZT sealant was studied on a 1 cm long full-thickness skin incision in male Sprague Dawley rats. The experiments consisted of 4 test articles: our ZT sealant as solo closure, or in combination with sutures (ZT sealant+suture) and two controls (sutures and Tisseel). The wound closure was studied at 7 and 14 days with 4 replicate rats per time point. Rats were anesthetized using isoflurane, and anesthetic maintenance from isoflurane in oxygen was given by mask. The hair from the dorsal section of each rat was removed and four equally spaced incisions were made per rat with a #15 blade until the underlying fascia was exposed. The suture closure was performed with two equally spaced buried intradermal sutures with an absorbable monofilament suture on a reverse cutting needle (4-0 Poliglecaprone 25, Monocryl PS-2, Ethicon). Approximately 50 μL of the ZT sealant or Tisseel was applied per incision to ensure that the incisions were filled. The rat remained anesthetized until the products in the non-sutured incisions fully dried. Instruments were re-sterilized between rats; a new blade and a new suture were used for each rat to minimize trauma.

During recovery, the rats were administered 0.65mg/kg buprenorphine extended-release (Ethiqa XR) for analgesia. The weight and activity levels of the rats were recorded initially and periodically during the experiment. The incision site was examined for signs of irritation, and photographs of the incisions were captured periodically to determine the wound closure rate. ImageJ was used to analyze the wound area from the photographs. At the end of each time point of the study, the rats were euthanized. The tissues from the injury site were harvested for further histopathological analysis. The ZT sealant+suture group was excluded from the final analysis due to inconclusive findings.

### Histopathological analysis

The tissue sections were fixed in formalin solution and embedded in paraffin. Four µm sections were cut and stained using Hematoxylin and Eosin (H&E) stain. The stained sections were observed and imaged using a DMLB microscope (Leica). Histomorphologic grading was conducted by a board-certified veterinary pathologist to analyze the sections. The grading scale was modified from a published assessment of the histological scoring system for cutaneous injury and is illustrated in Table S1[27,28] Low scores were the most healed, whereas high scores had the most pathology associated with them that preclude wound healing. Specifically, signs of epithelialization, inflammation, and fibrosis were assessed to confirm the remodeling of the dermis and epidermis at the end of each time point of the study.

### Statistics

Statistical analysis was performed using JMP software. When comparing only two groups, a two-tailed Student’s t-test or Dunnett’s test with equal variances was used with α = 0.05. When comparing more than two groups, analysis of variance (ANOVA) was used with α = 0.05. Normality was confirmed using the Shapiro-Wilk test, and equality of variance was validated using the Brown-Forsythe test. If the residuals of a comparison were non-normal or the variance was not equal, an appropriate transformation (e.g., Box-Cox, Johnson normalizing, log, or inverse transformation) was used. A Tukey post-hoc test was used to determine statistically significant groups. Statistical significance is indicated by letters or symbols such that groups that do not share the same letter are statistically different (p < 0.05).

## Results

### Design and formulation of sea squirt-inspired tissue sealants

We synthesized sea squirt-inspired gallol-functionalized proteins by crosslinking gallol- containing TA to the plant-based zein protein. These ZT sealants, composed of zein, TA, ethanol, and water, were combined in different ratios to yield tissue sealants. Zein is a protein byproduct obtained from corn as a result of ethanol extraction. Zein is extensively used for food packaging as adhesives and coatings.[29,30] Zein has been classified as “generally recognized as safe” (GRAS) by the United States Food and Drug Administration when used in those applications.[31,32] Moreover, zein has been extensively used for drug delivery applications in the literature.[33,34] Due to its amphiphilic nature, it is only partially soluble in water. Therefore, a combination of ethanol and water was used to dissolve zein. TA is a naturally occurring polyphenol and has been widely investigated for its antimicrobial, anti-inflammatory, and antitumor properties. Herein, TA was chosen as the source of gallol functional groups for sea squirt-inspired adhesion due to its wound-healing properties.[35–39] Our previous studies have shown that at neutral to basic pH, the gallols in TA oxidize and crosslink to various functional groups such as amines in the zein protein via Michael addition or radical–radical coupling (figure 1b).[23,24] This crosslinking mechanism is visible when the sealant turns from an amber color to a darker brown color within 24 h of synthesis. The sealant formulation with a higher concentration of TA (Z26T21) is visibly darker than the sealant formulations with a lower concentration of TA (Z40T7) as seen in figure 1c.

### Determination of best formulations using burst pressure test

The concentration of zein, TA, ethanol, and water for initial optimization studies was chosen based on our previous studies.[21–24] The zein concentrations were varied as 26 wt%, 30 wt%, and 40 wt%, and the TA concentrations were varied from 1-28 wt% for each of the zein concentrations. The TA concentration range studied for each zein concentration was limited by the final viscosity and consistency of the sealant. If the final consistency after mixing all the components was deemed to be too viscous (i.e., could not be pipetted) or had a water-like consistency, no further studies were carried out with those formulations.

Two formulations, 40 wt% zein with 7 wt% TA (Z40T7) and 26 wt% zein with 21 wt% TA (Z26T21) were chosen for further studies based on their overall highest burst pressures compared to all formulations (figure S1). To further improve the burst pressures of the sealants, TG was added to act as an interfacial crosslinker between the tissue and sealant. TG is a commercial food-grade enzyme that catalyzes the formation of isopeptide bonds between γ- carboxamide groups of glutamines and the ε-amino groups of lysines. [40,41] They have been widely used in literature to bond different proteins or proteins to non-protein molecules such as hyaluronic acid.[42–44] Herein, it was expected that TG would help bond proteins in zein to proteins in tissues. The burst pressure of Z26T21 did not show significant improvement with the introduction of TG since the mode of failure was cohesive (Figure 2a). This was not unexpected since Z26T21 also failed cohesively. However, using TG as a crosslinker led to a higher burst pressure of Z40T7. Similar to the results with only Z40T7, adhesive failure was observed for Z40T7+TG as well.

**Figure 2:**
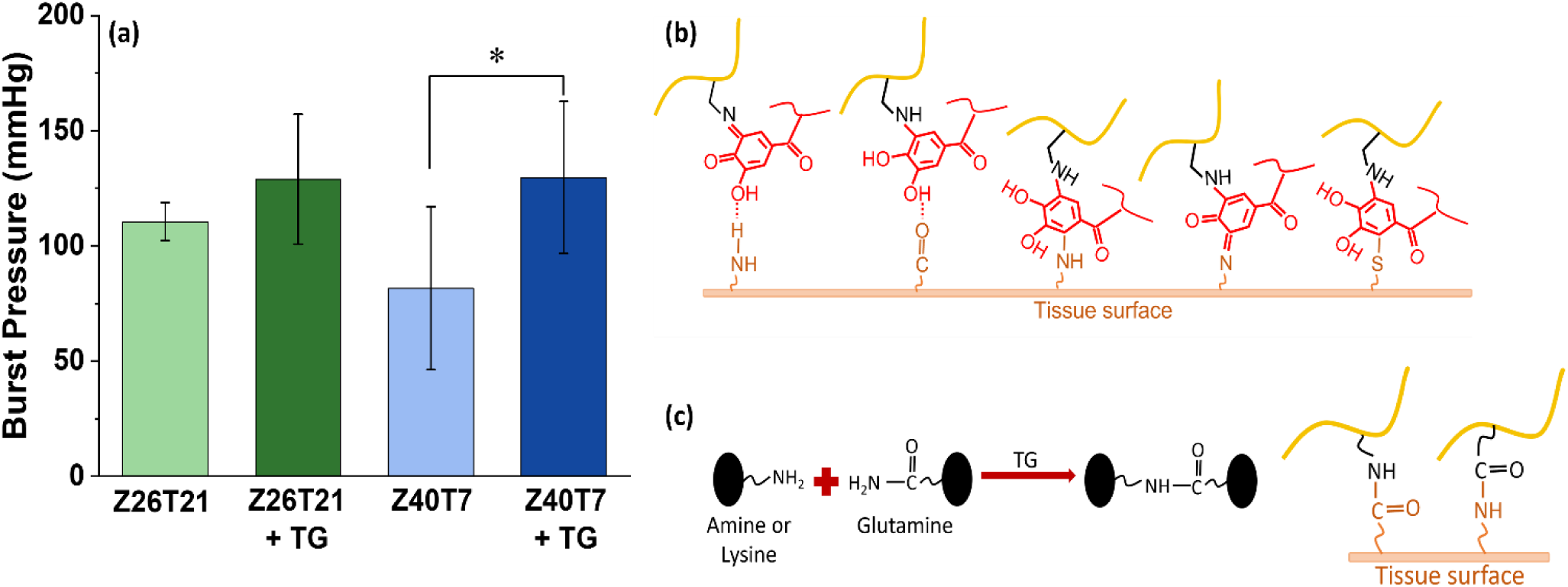
(a) The effect of using transglutaminase as an interfacial crosslinker. Different letters indicate distinct Tukey groups (p < 0.05). * indicates statistical significance with p < 0.05. Possible mechanism of adhesion of ZT sealants to tissues in the **(b)** absence of TG or **(c)** presence of TG.

The ZT sealants likely adhered to tissues via covalent and non-covalent interactions between gallols on the sealant and amines, carboxylic acids, imidazoles, and thiols on tissues (figure 2b). These tissue functional groups originate from amino acid residues in proteins and fatty acids in tissues.[2,45] The gallol groups in TA were oxidized to quinones which were able to react with functional groups on tissues via various pathways. Benzene ring of quinones undergo Michael addition reactions with amines, thiols, or imidazole on tissues. Quinones can form Schiff base linkages between carbonyl groups and amines.[2,45–47] Studies have shown that the majority of covalent interactions are based on Michael-type addition.[48] Moreover, physical crosslinking via hydrogen bonding could have contributed to quick adhesion. When TG was used as an interfacial crosslinker, it catalyzed the reaction either between the amine functional groups in zein and the glutamine residues on the tissue or between glutamine in zein and lysine in tissues (figure 2c).[42,49] Thus, TG acted as an interfacial crosslinker between zein and tissues to reinforce the adhesion strength.

### Burst pressures of ZT sealants on different tissue substrates

Initial optimization studies on porcine skin were conducted using a sealant volume of 20 μL. When the sealant volume was increased to 40 μL significant improvement in burst pressure was observed (figure 3a). Increasing the sealant volume above 40 μL did not affect the burst pressures, especially at a short cure time of 30 min because of incomplete curing. To find a targeted application for these ZT sealants, the adhesion performance of the sealants was studied and applied to eight other porcine tissue substrates including the stomach, sausage casing, small intestines, dura, liver, lungs, heart, and aorta (Figure 3). The choice of these tissue substrates was made after careful consideration to determine a potential, suitable application for the ZT sealant. Sausage casings were chosen as a representative tissue substrate similar to collagen sheets, which is a commonly studied material in the literature for assessing the adhesive properties of tissue adhesives.

**Figure 3:**
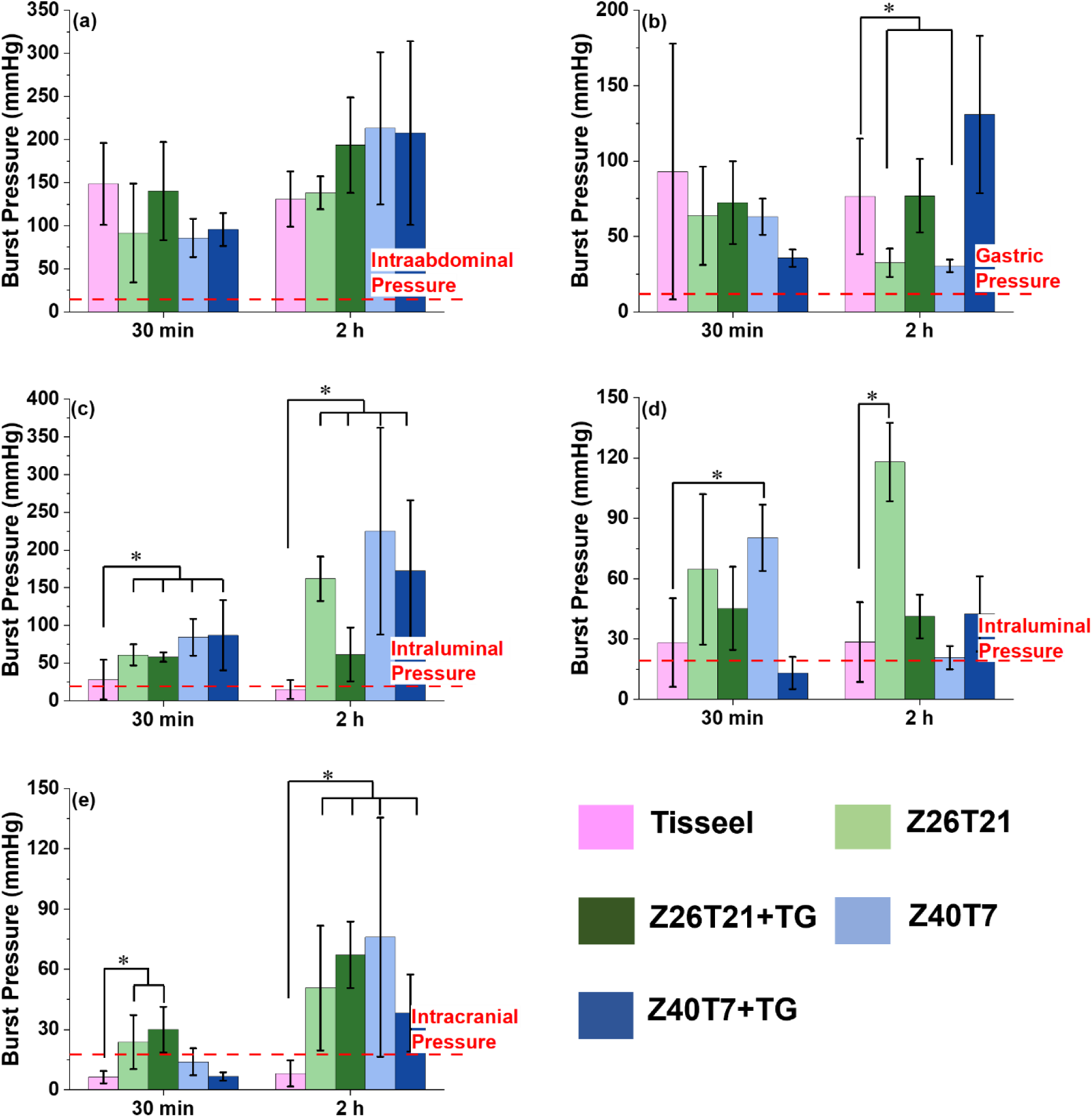
Burst pressures obtained for Tisseel, Z26T21, Z26T21+TG, Z40T7, and Z40T7+TG on porcine **(a)** sausage casing, **(b)** small intestine, **(c)** dura mater, **(d)** skin, and **(f)** stomach after a 30 min and 2 h cure. * indicates statistical significance compared to Tisseel with p < 0.05.

The performance of ZT sealants varied greatly on the different tissues. For instance, one of the highest burst pressure values (225 ± 137 mmHg) was recorded for sausage casings while the burst pressures obtained on lungs had some of the lowest values (11 ± 1.5 mmHg). Therefore, the obtained burst pressures were compared against normal physiological pressures experienced in each organ in humans. The burst pressures of ZT sealants exceeded physiological pressures on seven out of nine different tissues (all except the heart and aorta) studied here (figure 3 and S2).

The sealant performance was also benchmarked against Tisseel. The burst pressures of ZT sealants were similar to Tisseel for skin, stomach, lungs, and heart; however, its performance was much better than Tisseel for sausage casing, small intestine, dura, and liver (figure 3 and S2). Tisseel exhibited its best performance on aorta tissues for which its bursting pressures were higher than those for ZT sealants (figure 3d).

Each of the five sealants studied here (four ZT sealants and Tisseel) was best on different tissue substrates. Moreover, in most cases, the sealant that exhibited the highest burst pressure after 30 min, i.e., when the sealant was partially cured, was not the one that exhibited the highest burst pressure after completely curing the sealant at 2 h. Unexpectedly, in the case of some tissues (stomach, lungs, small intestines, and liver), the burst pressure after a 2 h cure was lower than that after a 30 min cure.

### *In vitro* cytocompatibility studies

Apart from having high adhesion strengths, a promising tissue sealant must also be cytocompatible to minimize toxicity when used within the body. Therefore, the cytocompatibility of the leachates from ZT sealants was studied at different leachate dilutions using cell viability and cell metabolic activity assays. From figures 4a and 4b, Z26T21 exhibited viable cells when the leachate was diluted to 10x and 100x, whereas Z40T7 showed viable cells at all leachate concentrations. Moreover, the metabolic activity assay in figure 4c and 4d showed that cell metabolic activity increased for both Z26T21 and Z40T7 with increasing leachate dilution. There was no difference in metabolic activity between the positive control and groups with Z26T21 diluted to 100x and Z40T7 diluted to 10x and 100x. However, undiluted Z26T21 had statistically similar cell metabolic activity to the negative control indicating slight leachate toxicity for Z26T21 under the studied conditions.

**Figure 4:**
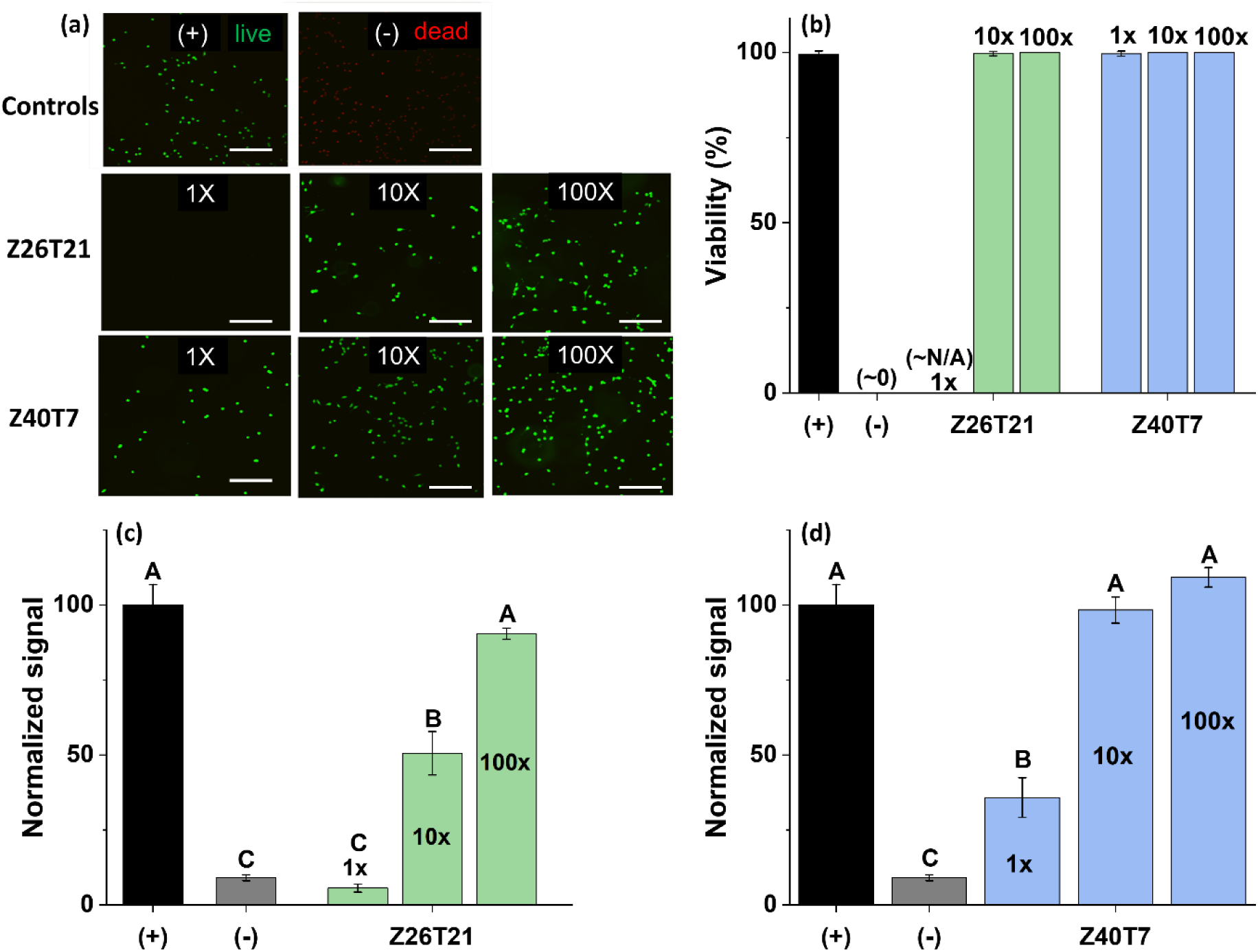
Cell viability **(a)** images and **(b)** results for cells exposed to leachate at 1x, 10x, and 100x dilution. The scale bars represent 300 µm. Metabolic activity of cells exposed to leachate from **(c)** Z26T21 and **(d)** Z40T7 at different dilutions. Different letters indicate distinct Tukey groups (p < 0.05).

The slight cytotoxicity at high concentrations of TA (Z26T21) could be attributed to the high acidity from TA as well as the autoxidation of gallol in TA into its quinone form and subsequent generation of reactive oxygen species (ROS) such as hydrogen peroxide and superoxide anion.[50,51] These oxidizing agents can negatively interact with cells due to their high reactivity.[52]

### Ex vivo studies

The ZT sealants exhibited adhesion strengths higher than physiological pressures on most of the tissue substrates studied here. However, to determine their practical applicability, the *in vitro* results were validated using *ex vivo* models. Moreover, during *in vitro* studies, the sealants were cured on tissues for as long as 30 min and 2 h which may not be practically applicable when surgeons need to close actively leaking wounds. Therefore, two *ex vivo* models were developed based on the two most promising tissue substrates.

An *ex vivo* stomach model was designed to study the tissue sealing ability of the sealants on an unpressurized leak such as those arising from gastrointestinal perforations or minor anastomotic leaks. As shown in figure 5a, when a 3 mm puncture was introduced onto the stomach tissue, dyed water leaked from the puncture. An instant seal was formed when sealing the puncture with Z40T7 and Z26T21 underwater, and no further leak was detected.

**Figure 5:**
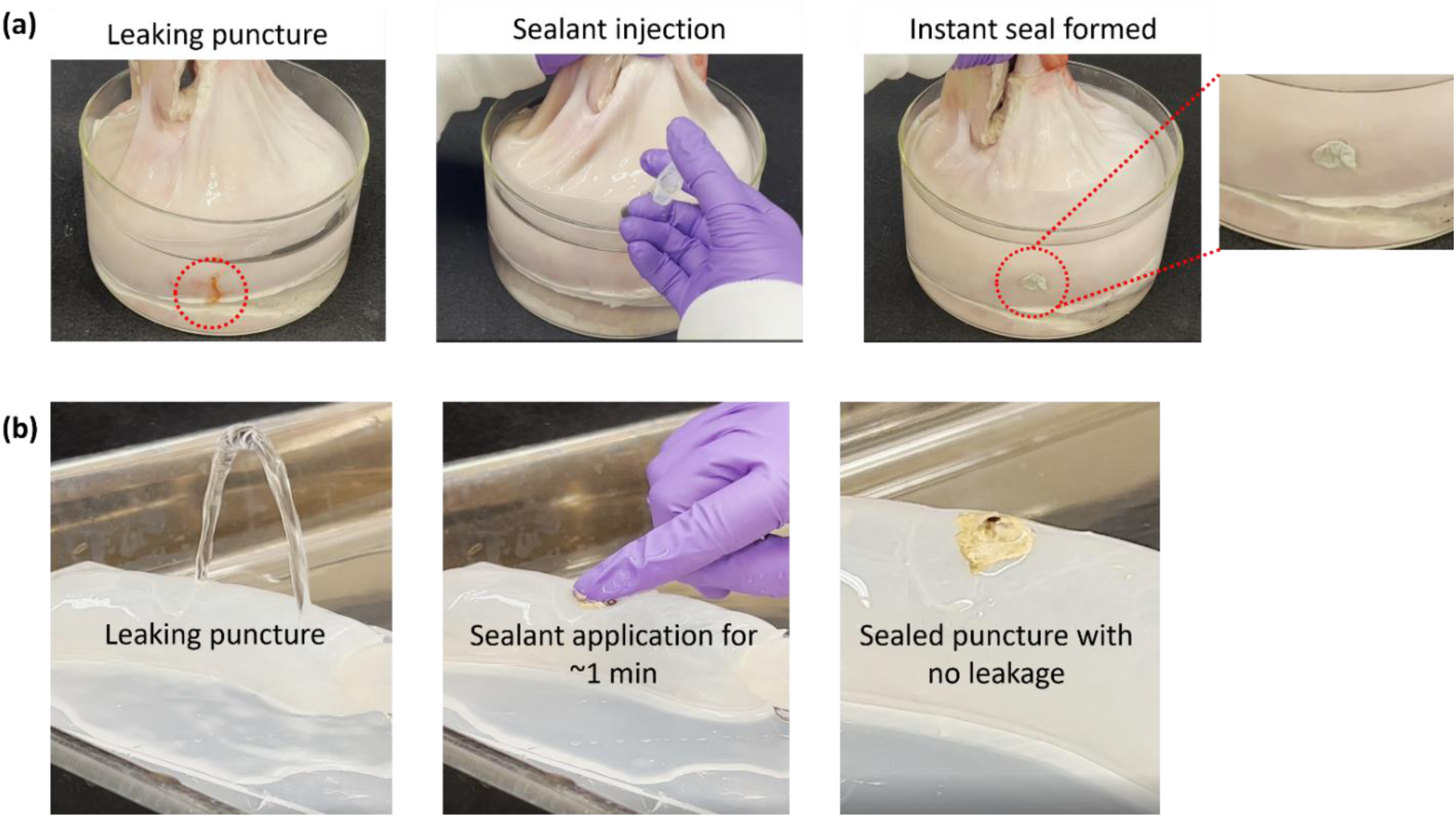
(a) *Ex vivo* model of stomach tissue showing the instant sealing of a leaking puncture underwater using ZT sealants. **(b)** *Ex vivo* model of sausage casings showing the sealing of leaking puncture in less than 1 min using ZT sealants at pressures higher than normal intraluminal pressures.

Similarly, the ability of the sealants to seal actively leaking punctures under pressurized conditions was also tested using an *ex vivo* sausage casing model. This model was studied to validate the ability of ZT sealants to seal tissues in situations such as hemorrhaging from a gunshot wound or trauma. Water flowed continuously through the sausage casing at a pressure of 30 mmHg (higher than normal physiological pressure of 20 mmHg in intestines). As shown in figure 5b, Z40T7 was applied to a leaking puncture for 30-40 s using mild pressure. The puncture instantly stopped leaking. The seal remained intact after 1 h. Similarly, Z26T21 also sealed the puncture within less than a minute. The tissue sealing ability could not be studied using TG for either of the models since it instantly dissolved in water.

### *In vivo* wound healing studies

The promising results obtained from *in vitro* and *ex vivo* studies were further validated using an *in vivo* wound healing model in rodents. Since Z40T7 showed less toxicity in cytocompatibility studies, this sealant was chosen for this study. ZT sealant was applied to full-thickness skin incisions in rats and compared against two controls, Tisseel and sutures. The rats were studied for two different time points of 7 and 14 days. From figure 6a, the suture group showed complete wound healing by day 10, whereas the Z40T7 group healed completely by day 14. In the case of the Tisseel group, the wounds had healed for all mice except for one where there was still some scabbing left on day 14. There were no statistical differences between the wound closure rates of Tisseel and Z40T7 as shown in figure 6b. The wound closure rate for the suture group was not calculated due to the absence of an open wound.

**Figure 6:**
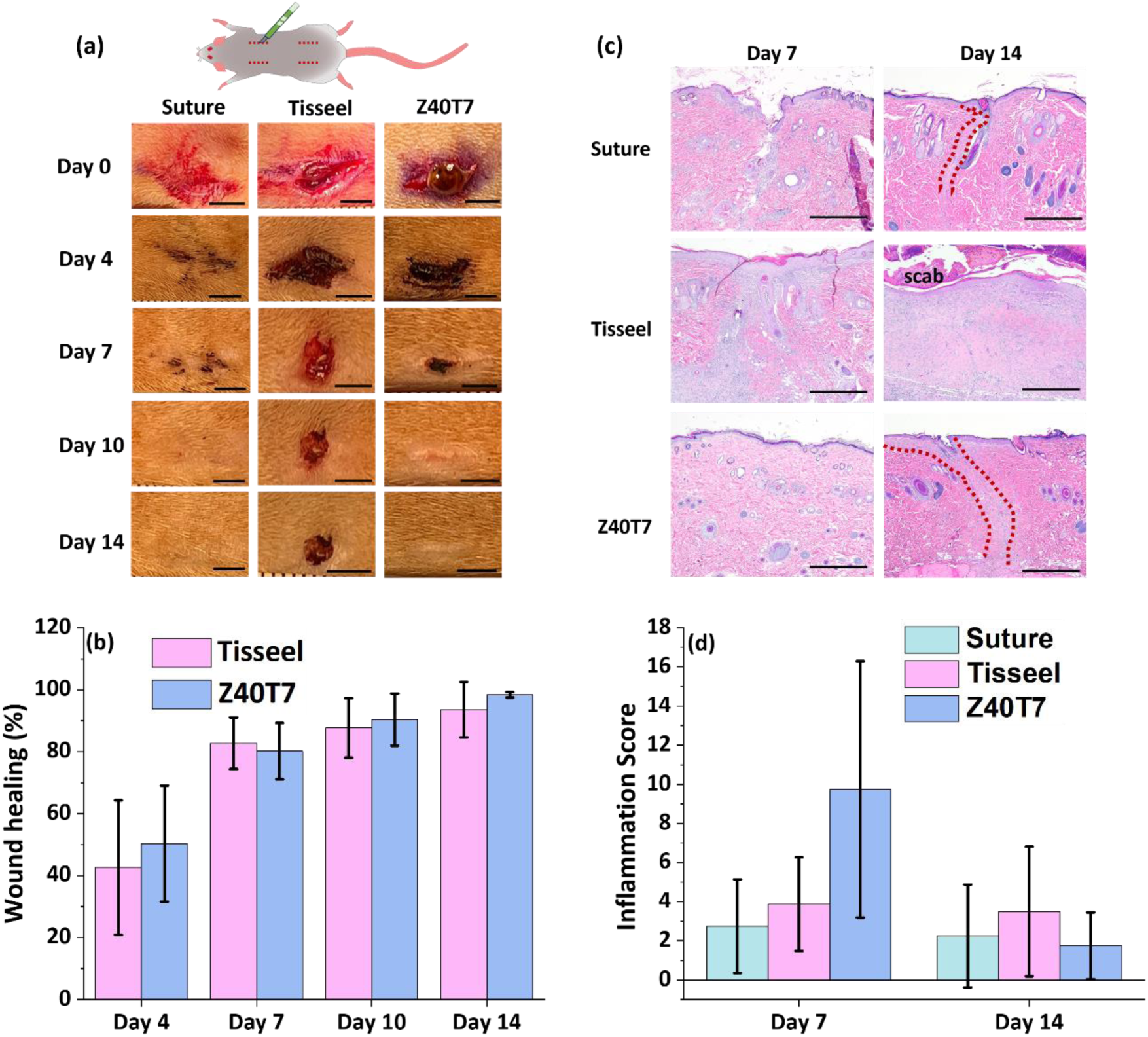
(a) Photographs of wound area at days 0 to 14 for suture, Tisseel, and Z40T7. The scale bars represent 4 mm. **(b)** Graph comparing the wound closure rate of Z40T7 to that of Tisseel. **(c)** H&E stained wound areas for suture, Tisseel, and Z40T7 on days 7 and 14. The scale bars represent 500 μm. The wound area is marked in red. **(d)** Graph comparing the overall inflammation score of Z40T7 to that of Tisseel and sutures at days 7 and 14.

Further, histopathological evaluation of all the wound sites was carried out for both the 7-day and 14-day cohorts via H&E staining (figure S5, S6, and 6c). Three of the four rats in the 7- day cohort treated with Z40T7 had marked inflammation including multinucleated giant cells and little to no signs of healing. These observations are consistent with the occurrence of a foreign body reaction which is typical in the case of implantation of a foreign material. The suture and Tisseel groups were similar to each other had mild to moderate changes in inflammation and showed evidence of appropriate wound healing. On day 7, the wound site treated with Z40T7 is possibly more inflamed although the inflammation scores were not found to be statistically different from the Tisseel or the suture group (figure 6d and S7). Moreover, the incorporation of catechols or gallols has been linked to the formation of stronger foreign body reactions due to ROS generation in some studies.[53] However, on day 14, the inflammation observed for Z40T7 was similar when compared to all wound closure methods studied here (figure 6d and S8). The inflammation observed with the application of Z40T7 had completely subsided within 14 days. All the groups showed complete re-epithelization with keratinization and evidence of effective wound bed closure and re-modeling.

## Conclusions and Future Perspectives

We developed a completely bio-based, easily synthesizable, and low-cost sea squirt- inspired tissue sealant based on zein (corn protein) and TA. Two sealant formulations (Z26T21 and Z40T7) were identified by varying the ratio of zein to tannic acid and studying their burst pressures on porcine skin. A food-grade interfacial crosslinker, TG, enhanced the burst pressures of the sealants through additional crosslinking with the tissues. The four ZT sealant formulations (Z26T21, Z26T21+TG, Z40T7, and Z40T7+TG) were studied on nine different tissues at short and long cure times. The sealant’s performance was also benchmarked against the FDA-approved sealant Tisseel. Our studies showed that the performance of all the sealants studied here varied when studied on different tissues. Each of the sealants studied here showed their best performance on a different tissue substrate. Our findings suggest that adhesion between a sealant and tissue of interest is a complex phenomenon and is affected both by the composition and chemistry of the tissue and the sealant. The matching of mechanical properties such as elastic modulus or stiffness of the sealant with underlying tissues is an essential consideration.[46] The mechanical disparity between the sealants and the underlying tissue can lead to stresses concentrated at the tissue−sealant interface which could disrupt their chemical and physical links and lead to delamination or failure.[2] The modulus of the sealant is affected by its composition and cure time. For the same composition, the sealant can become stiffer after completely curing, which can lead to a mismatch of modulus between tissue and sealant. The exact reason as to why certain sealants perform well only on certain tissues is not clear and will need further probing. These studies strengthen the need to develop tissue adhesives and sealants that are specifically tailored to a particular tissue type to ensure complete wound closure until tissue regeneration takes place.

The burst pressures of the ZT sealants exceeded physiological pressures in seven out of the nine tissues studied here. The sealant performance was better than or on par with the performance of Tisseel on eight out of the nine tissues. When tested for their practical applicability in *ex vivo* models of sausage casing and stomach tissues, instant wound closure was achieved in less than a minute. The sealants were cytocompatible and supported complete wound healing *in vivo* when applied on skin incisions in rats with minimal inflammation. The observed wound healing was also comparable to sutures and Tisseel. With these promising results, we envision the application of our low-cost sea squirt-inspired tissue sealants to seal both topical wounds as well as internal wounds in future surgical settings. Moreover, this study not only describes the application of sea squirt-inspired sealants for wound closure but also demonstrates the need for careful material selection and design for targeted end application.

## Acknowledgments

This project was funded with support from the Indiana Clinical and Translational Sciences Institute, which is funded in part by Award Number UM1TR004402 from the National Institutes of Health, National Center for Advancing Translational Sciences, Clinical and Translational Sciences Award (A.V.M). J.C.L., V.R.R., and A.D.C. acknowledge funding from the pilot project program through the Engineering in Medicine Institute. The authors also acknowledge funding from the Purdue Davidson School of Chemical Engineering (J.C.L. and A.V.M.), National Science Foundation (grants DMR-2104783 to J.C.L. and J.J.W. and a Graduate Fellowship grant DGE- 1842166 to J.E.T.), Office of Naval Research (grants N000-14-19-1-2342 and N000-14-22-1-2408 to J.J.W.), and a Lillian Gilbreth Postdoctoral Fellowship from Purdue University (A.V.M.). This work was completed in collaboration with the Pre-Clinical Research Laboratory and the Histology Research Laboratory, which are both part of the Center for Comparative Translational Research (Purdue University, College of Veterinary Medicine). Robyn McCain provided help and guidance with experiments.

## References

[1] P. Heher, J. Ferguson, H. Redl, P. Slezak, Expert Rev Med Devices 2018, 15, 747.

[2] S. Nam, D. Mooney, Chem Rev 2021, 121, 11336.

[3] N. Gillman, D. Lloyd, R. Bindra, R. Ruan, M. Zheng, Expert Rev Med Devices 2020, 17, 443.

[4] Z. Bao, M. Gao, Y. Sun, R. Nian, M. Xian, Mater Sci Eng C 2020, 111, 110796.

[5] S. Tarafder, G. Y. Park, J. Felix, C. H. Lee, Acta Biomater 2020, 117, 77.

[6] H. Hill, J. F. B. Chick, A. Hage, R. N. Srinivasa, Diagn Interv Radiol 2018, 24, 98.

7. J. E. Torres, S. Hollingshead, D. Boucher, J. C. Liu, Biomimetic Protein Based Elastomers: Emerging Materials for the Future, Royal Society of Chemistry 2022, 10, 210.

[8] E. Borie, E. Rosas, G. Kuramochi, S. Etcheberry, S. Olate, B. Weber, Biomed Res Int 2019, 2019, DOI 10.1155/2019/8217602.

[9] M. Beudert, M. Gutmann, T. Lühmann, L. Meinel, ACS Biomater Sci Eng 2022, 8, 2220.

[10] D. W. R. Balkenende, S. M. Winkler, P. B. Messersmith, Eur Polym J 2019, 116, 134.

[11] L. Ge, S. Chen, Polymers 2020, 12, 939.

[12] Q. Guo, J. Chen, J. Wang, H. Zeng, J. Yu, Nanoscale 2020, 12, 1307.

[13] A. Menezes, H. Douglas Melo Coutinho, A. Gonçalves Wanderley, J. Ribeiro-Filho, J. Melrose, Molecules 2022, 27, 8982.

14. Y. Rao, G. Wan, Adv Struct Adhes Bonding 2023, 953.

[15] C. Heinritz, X. J. Ng, T. Scheibel, Adv Funct Mater 2023, 2303609.

[16] J. Delroisse, V. Kang, A. Gouveneaux, R. Santos, P. Flammang, Convergent Evolution: Animal Form and Function, Cham: Springer International Publishing 2023, 523.

[17] K. Zhan, C. Kim, K. Sung, H. Ejima, N. Yoshie, Biomacromolecules 2017, 18, 2959.

[18] M. Cai, M. Sugumaran, W. E. Robinson, Comp Biochem Physiol B Biochem Mol Biol 2008 151, 110.

[19] S. W. Taylor, B. Kammerer, E. Bayer, Chem Rev 1997, 97, 333.

[20] S. W. Taylor, M. M. Ross, J. Herbert Waite, Arch Biochem Biophys 1995, 324, 228.

[21] G. Schmidt, J. T. Woods, L. X. B. Fung, C. J. Gilpin, B. R. Hamaker, J. J. Wilker, Adv Sustain Syst 2019, 3, 1900077.

[22] G. Schmidt, K. H. Smith, L. J. Miles, C. K. Gettelfinger, J. A. Hawthorne, E. C. Fruzyna, J. J. Wilker, Adv Sustain Syst 2022, 6, 2100392.

[23] G. Schmidt, B. R. Hamaker, J. J. Wilker, Adv Sustain Syst 2018, 2, 1700159.

[24] G. Schmidt, P. E. Christ, P. E. Kertes, R. V. Fisher, L. J. Miles, J. J. Wilker, ACS Appl Mater Interfaces 2023, 15, 32863.

[25] ASTM F2392-04- Standard Test Method for Burst Strength of Surgical Sealants, https://www.astm.org/standards/f2392.

26. ISO 10993-5:2009 - Biological evaluation of medical devices — Part 5: Tests for in vitro cytotoxicity,” https://www.iso.org/standard/36406.html.

[27] G. S. Lazarus, D. M. Cooper, D. R. Knighton, D. J. Margolis, R. E. Percoraro, G. Rodeheaver, M. C. Robson, Wound Repair Regen 1994, 2, 165.

[28] V. Lomash, S. E. Jadhav, F. Ahmed, R. Vijayaraghavan, S. C. Pant, Hum Exp Toxicol 2012, 31, 588.

[29] F. Garavand, D. Khodaei, N. Mahmud, J. Islam, I. Khan, S. Jafarzadeh, R. Tahergorabi, I. Cacciotti, Crit Rev Food Sci Nutr 2022, 12, 1.

[30] M. R. Kasaai, Trends Food Sci Technol 2018, 79, 184.

31. Federal Register: Zein; Exemption From the Requirement of a Tolerance, https://www.federalregister.gov/documents/2023/02/27/2023-03831/zein-exemption-from-the-requirement-of-a-tolerance.

32. CFR - Code of Federal Regulations Title 21, https://www.accessdata.fda.gov/scripts/cdrh/cfdocs/cfcfr/CFRSearch.cfm?fr=184.1984.

[33] K. H. Bae, I. De Marco, Polymers 2022, 14, 2172.

34. C. J. Pérez-Guzmán, R. Castro-Muñoz, Processes 2020, 8, 1376.

[35] H. Jafari, P. Ghaffari-Bohlouli, S. V. Niknezhad, A. Abedi, Z. Izadifar, R. Mohammadinejad, R. S. Varma, A. Shavandi, J Mater Chem B 2022, 10, 5873.

[36] Z. Guo, W. Xie, J. Lu, X. Guo, J. Xu, W. Xu, Y. Chi, N. Takuya, H. Wu, L. Zhao, J Mater Chem B 2021, 9, 4098.

[37] B. Kaczmarek, Materials 2020, 13, 3224.

[38] C. Chen, H. Yang, X. Yang, Q. Ma, RSC Adv 2022, 12, 7689.

[39] A. Baldwin, B. W. Booth, J Biomater Appl 2022, 36, 1503.

[40] K. Yokoyama, N. Nio, Y. Kikuchi, Appl Microbiol Biotechnol 2004, 64, 447.

[41] M. Kieliszek, A. Misiewicz, Folia Microbiol (Praha) 2014, 59, 241.

[42] M. K. McDermott, T. Chen, C. M. Williams, K. M. Markley, G. F. Payne, Biomacromolecules 2004, 5, 1270.

[43] J. G. Fernandez, S. Seetharam, C. Ding, J. Feliz, E. Doherty, D. E. Ingber, Tissue Eng Part A 2017, 23, 135.

[44] C. V. L. Giosafatto, A. Fusco, A. Al-Asmar, L. Mariniello, Int J Mol Sci 2020, 21, 3656.

[45] H. A. Lee, E. Park, H. Lee, H. A. Lee, E. Park, H. Lee, Adv Mater 2020, 32, 1907505.

46. G. M. Taboada, K. Yang, M. J. N. Pereira, S. S. Liu, Y. Hu, J. M. Karp, N. Artzi, Y. Lee, Nat Rev Mater 2020, 5, 310.

[47] J. Yang, M. A. Cohen Stuart, M. Kamperman, Chem Soc Rev 2014, 43, 8271.

[48] J. Yang, V. Saggiomo, A. H. Velders, M. A. C. Stuart, M. Kamperman, *PLoS One* 2016, 11, e0166490.

[49] J. G. Fernandez, S. Seetharam, C. Ding, J. Feliz, E. Doherty, D. E. Ingber, Tissue Eng Part A 2017, 23, 135.

50. D. T. Sawyer, J. S. Valentine, Acc. Chem. Res 1981, 14, 393.

[51] M. Mochizuki, S. I. Yamazaki, K. Kano, T. Ikeda, Biochim Biophys Acta - Gen Subj 2002, 1569, 35.

[52] S. Hollingshead, H. Siebert, J. J. Wilker, J. C. Liu, J Biomed Mater Res A 2022, 110, 43.

[53] H. Meng, Y. Li, M. Faust, S. Konst, B. P. Lee, Acta Biomater 2015, 17, 160.

