## Supplemental information for "Sea squirt-inspired bio-derived tissue sealants"

### (Supplementary Information)

Aishwarya V. Menon[1,2] , Jessica E. Torres[1], Abigail D. Cox[3], Marije Risselada[4], Gudrun Schmidt[2], Jonathan J. Wilker[2,5#,*], Julie C. Liu[1,6,#,*]

*1. Davidson School of Chemical Engineering, Purdue University, West Lafayette, IN 47907,* USA

*2. Department of Chemistry, Purdue University, West Lafayette, IN 47907, USA*

*3. Department of Comparative Pathobiology, Purdue University, West Lafayette, IN 47907, USA*

*4. Department of Veterinary Clinical Sciences, Purdue University, West Lafayette, IN 47907, USA*

*5. School of Materials Engineering, Purdue University, West Lafayette, IN 47907, USA*

*6. Weldon School of Biomedical Engineering, Purdue University, West Lafayette, IN 47907, USA*

*[#] Contributed equally as co-senior authors*

** Corresponding author*

**Table S1:** Scale used for histomorphologic grading of wound sites.

| Scale | Epithelization | Inflammation (PMN) | Macrophages (multinucleated) | Granulation tissue maturation | Amount of granulation tissue |
|---|---|---|---|---|---|
| 0 | Keratinization | Minimal | Absent | Organized fibroblasts with collagen and fewer vessels | Normal dermis, minimal granulation tissue remains |
| 1 | Bridging of cells | Mild | Mild | Fibroblasts and ECM forming layers, vessels oriented perpendicular | Thick granulation over the whole wound bed |
| 2 | Migration of cells | Moderate | Moderate | Macrophages and fibroblasts loosely arranged, few vessels | Wound bed partially filled with granulation tissue |
| 3 | Absent | Marked | Marked | Absent | Absent |

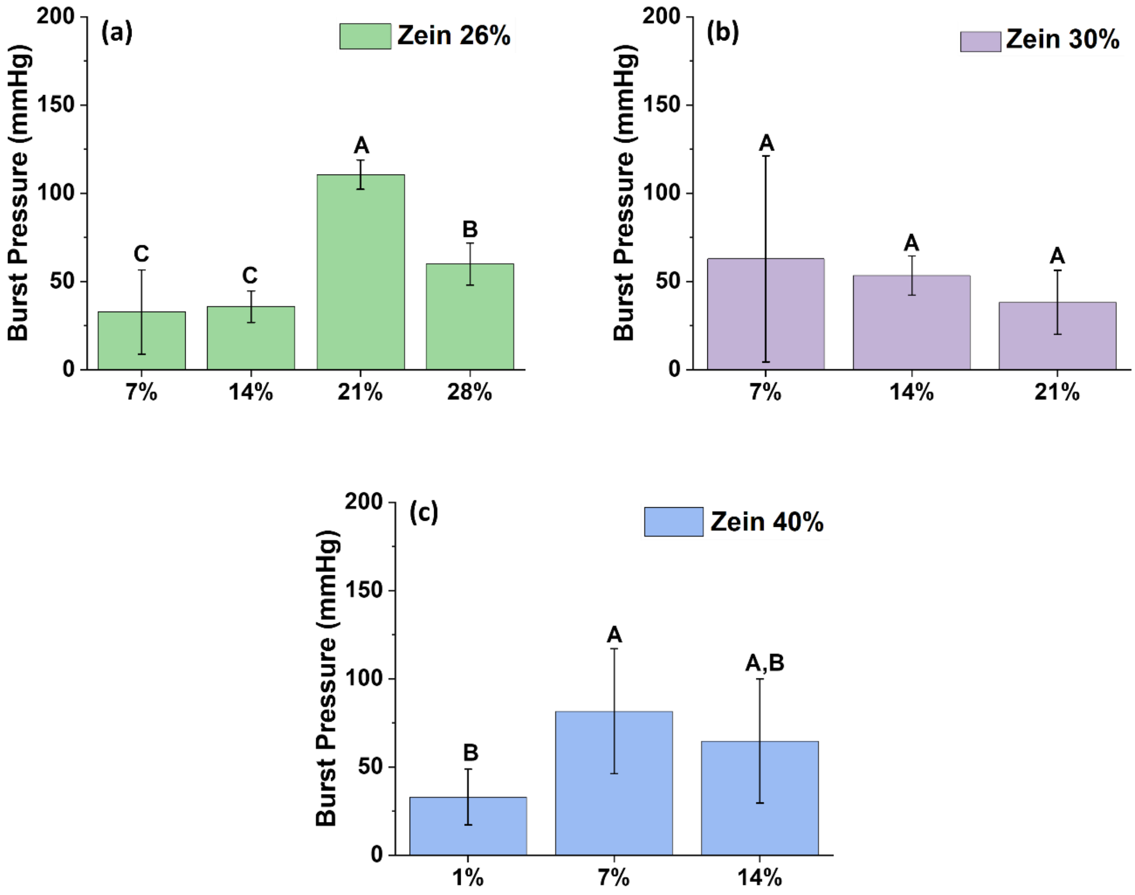


**Figure S1:** Effect of varying TA concentration from **(a)** 7-28 wt.% at 26 wt.% zein **(b)** 7-21 wt.% at 30 wt.% zein **(c)** 1-14 wt.% at 40 wt.% zein.

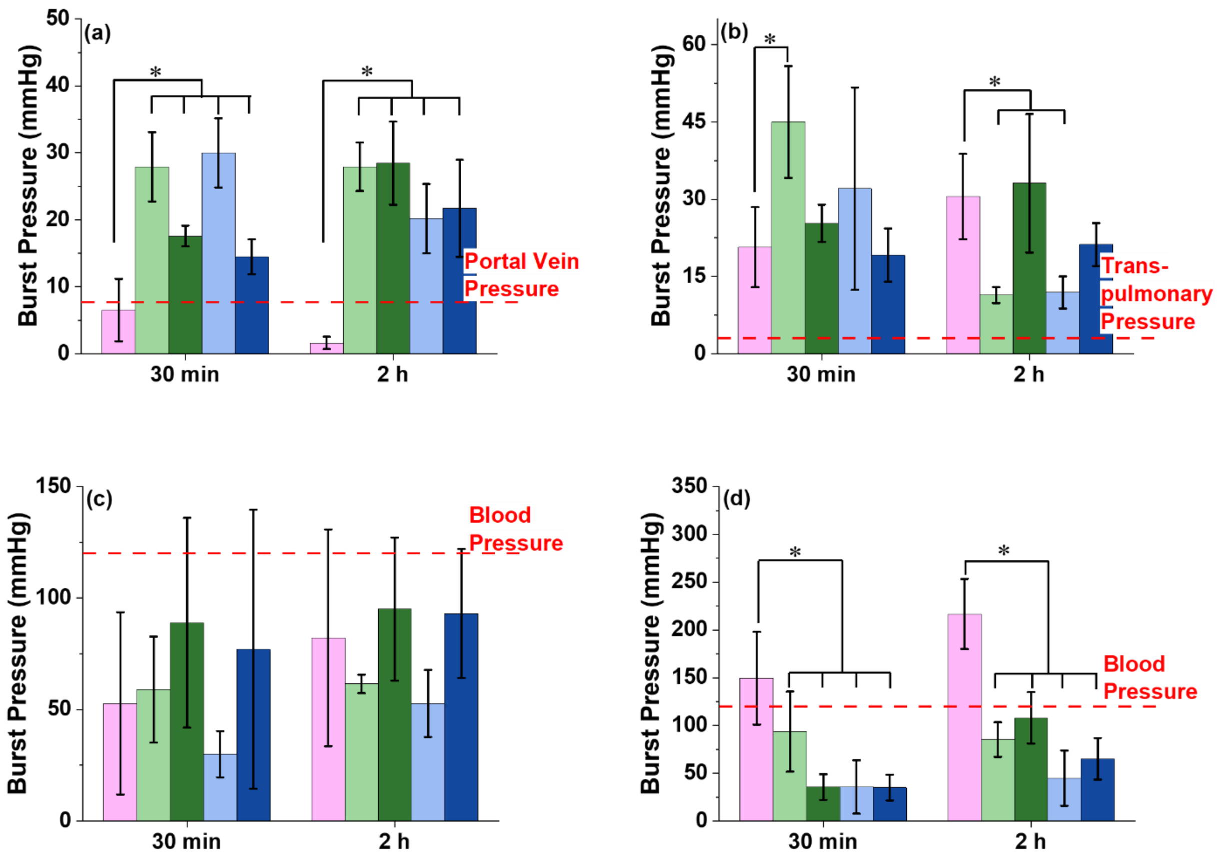


**Figure S2:** Burst pressures obtained for Tisseel, Z26T21, Z26T21+TG, Z40T7, and Z40T7+TG on porcine (a) liver (b) lungs (c) heart (d) aorta at 30 min and 2 h cure times. * indicates statistical significance with $p < 0.05$.

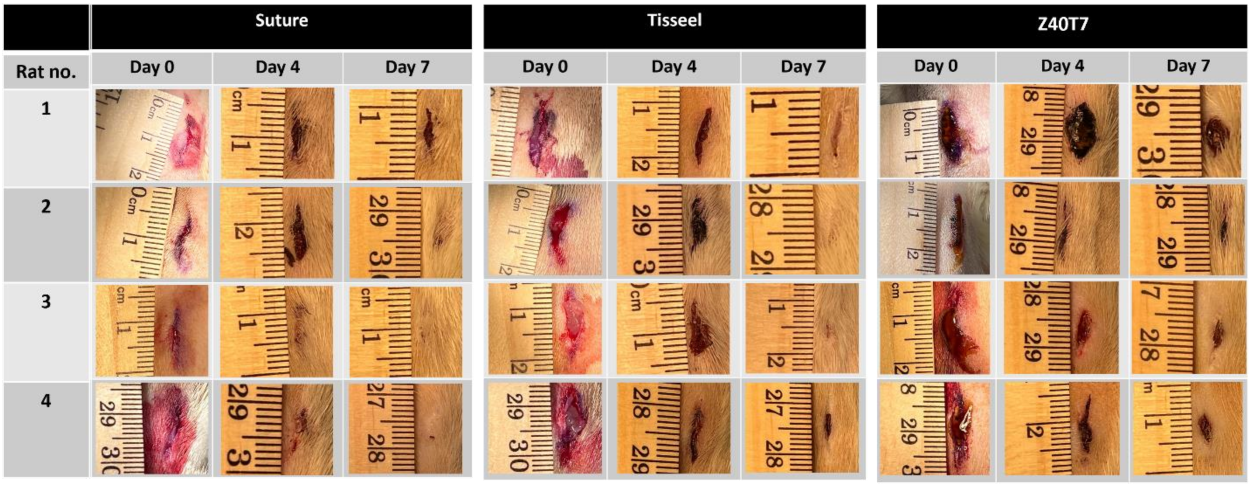


**Figure S3:** Photographs of skin incisions closed using suture, Tisseel, and Z40T7 for the 7-day cohort.

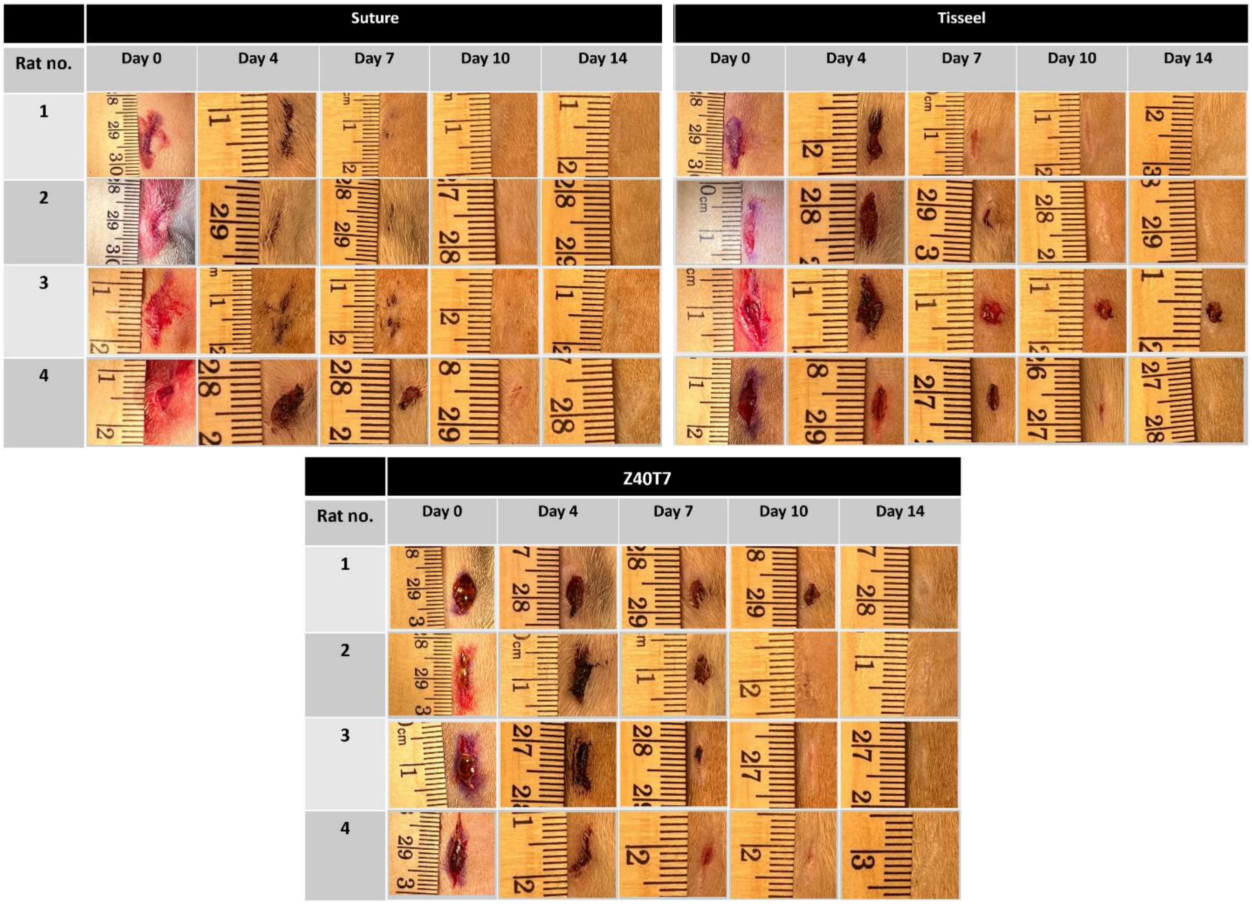


**Figure S4:** Photographs of skin incisions closed using suture, Tisseel, and Z40T7 for the 14-day cohort.

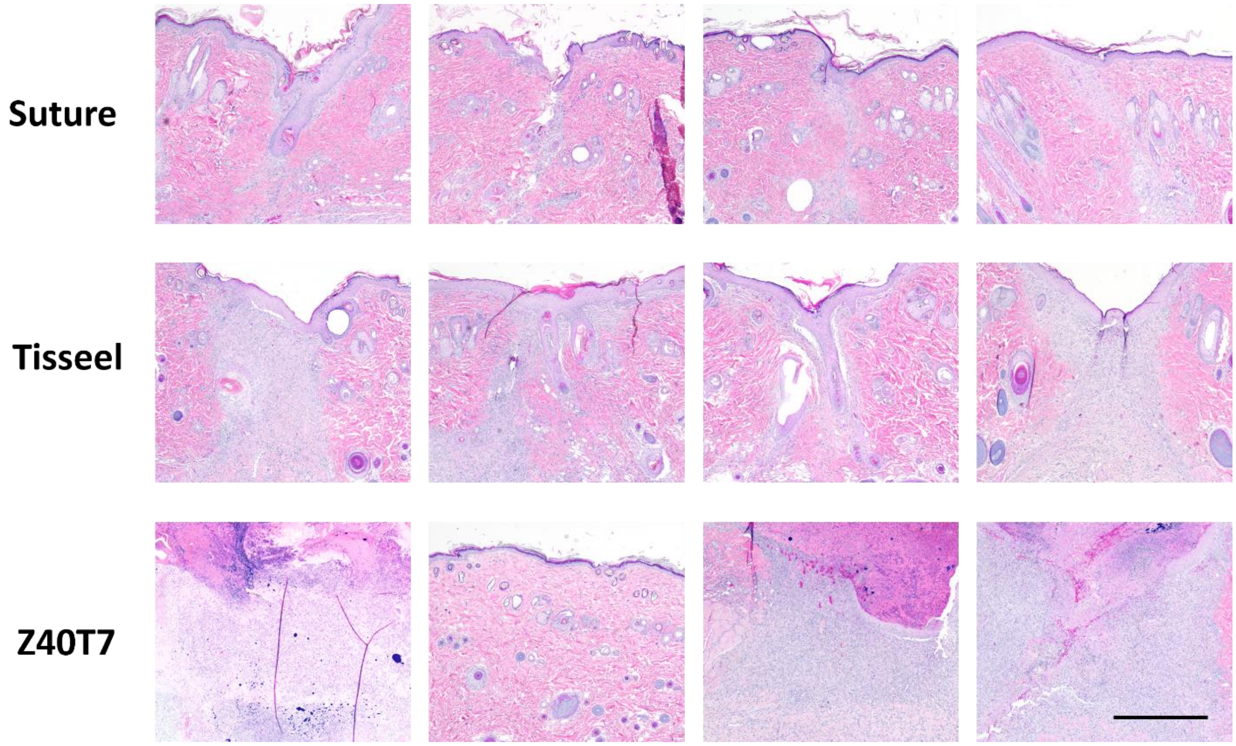


**Figure S5:** Micrographs of H&E stained skin incisions closed using suture, Tisseel, and Z40T7 for the 7-day cohort. The scale bar represents 500 µm.

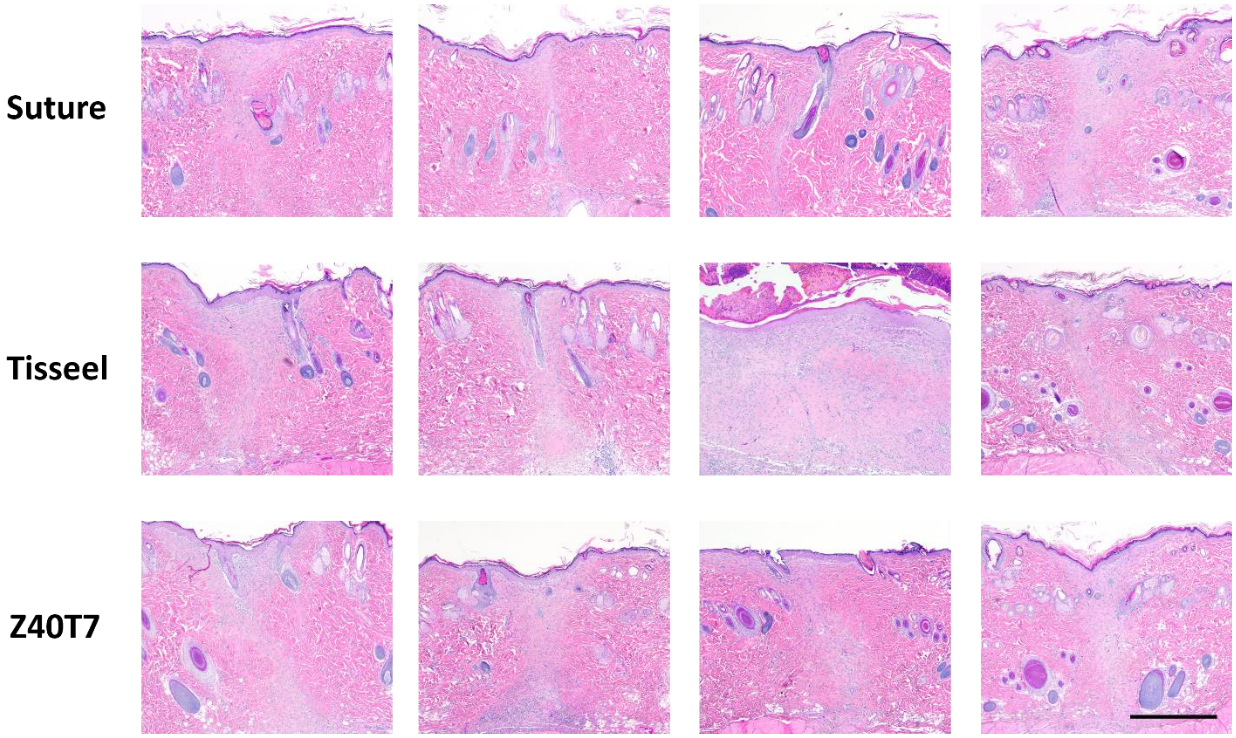


**Figure S6:** Micrographs of H&E stained skin incisions closed using suture, Tisseel, and Z40T7 for the 14-day cohort. The scale bar represents 500 µm.

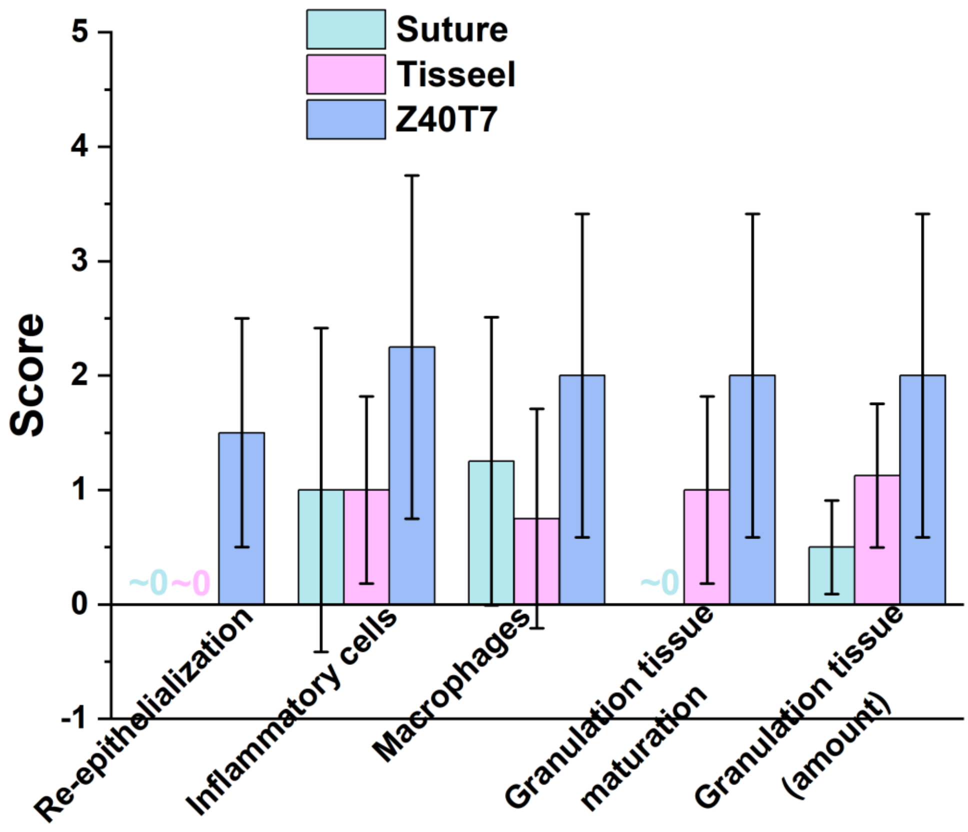


**Figure S7:** Average of the scores of different histomorphological observations used to assess skin incisions closed using suture, Tisseel, and Z40T7 in the 7-day cohort. Low scores are the most healed. High scores have the most pathology associated with them that will preclude wound healing.

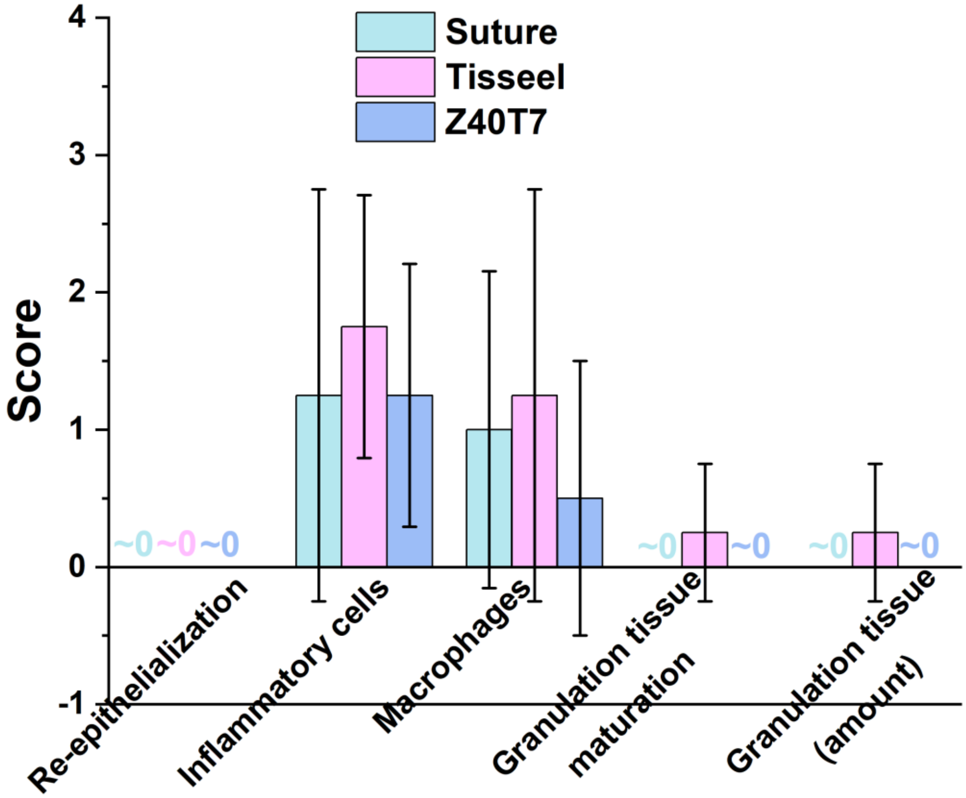


**Figure S8:** Average of the scores of different histomorphological observations used to assess skin incisions closed using suture, Tisseel, and Z40T7 in the 14-day cohort. Low scores are the most healed. High scores have the most pathology associated with them that will preclude wound healing.
